# *De novo* β-alanine synthesis in α-proteobacteria involves a β-alanine synthase from the uracil degradation pathway

**DOI:** 10.1101/663849

**Authors:** Mariana López-Sámano, Luis Fernando Lozano-Aguirre Beltrán, Rosina Sánchez-Thomas, Araceli Dávalos, Tomás Villaseñor, Jorge Donato García-García, Alejandro García-de los Santos

## Abstract

β-alanine synthesis in bacteria occurs by the decarboxylation of L-aspartate as part of the pantothenate synthesis pathway. In the other two domains of life we find different pathways for β-alanine formation, such as sources from spermine in plants, uracil in yeast and by transamination reactions in insects and mammals. There are also promiscuous decarboxylases that can decarboxylate aspartate. Several bioinformatics studies about the conservation of pantothenate synthesis pathway performed on bacteria, archaea and eukaryotes, have shown a partial conservation of the pathway. As a part of our work, we performed an analysis of the prevalence of reported β-alanine synthesis pathways in 204 genomes of alpha-proteobacteria, with a focus on the *Rhizobiales* order. The aim of this work was to determine the enzyme or pathway used to synthetize β-alanine in *Rhizobium etli* CFN42. Our bioinformatics analysis showed that this strain encodes the pyrimidine degradation pathway in its genome. We obtained a β-alanine synthase (*amaB)* mutant that was a β-alanine auxotroph. Complementation with the cloned gene restored the wild type phenotype. Biochemical analysis confirmed that the recombinant AmaB catalyzed the formation of β-alanine from 3-Ureidopropionic acid *in vitro*. Here we show a different way in bacteria to produce this essential metabolite.

**Importance:** Since the pioneer studies of Cronan (1980) on β-alanine synthesis in *E. coli*, it has been assumed that the decarboxilation of aspartate by the L-aspartate-α-decarboxylase it’s the main enzymatic reaction for β-alanine synthesis in bacteria. Forty years later, while we were studying the pantothenic acid synthesis in rhizobia, we demonstrate that a numerous and diverse group of bacteria classified as α-proteobacteria synthesize β-alanine *de novo* using β-alanine synthase, the last enzyme from the reductive pathway for uracil degradation.

Additionally, there is a growing interest in β-amino acid due to its remarkable pharmaceuticals properties as hypoglycemic, antiketogenic and anti-fungal agents.

## Introduction

β-alanine is a non-proteinogenic β-amino acid naturally occurring in all living organisms. In prokaryotes β-ala is indispensable for the synthesis of pantothenate, the precursor of the essential cofactor coenzyme A (CoA). CoA is the source of 4’-phosphopantetheine for fatty acid and polyketides synthesis (1). In eukaryotes, β-amino acids and β-peptides play important roles in the regulation of nutritional metabolism, immunity and the central nervous system (2).

The major pathway for β-ala synthesis in *Escherichia coli* is the decarboxylation of aspartate by aspartate decarboxylase (3). The protein ADC is a pyruvoyl-dependent enzyme that is initially synthesized as a zymogen (Pro-ADC). A cleavage of pro-ADC occurs between Gly24 and Ser25, creating the active-site pyruvoyl moiety. Stuecker (4) proposed two classes of ADC based on the type of cleavage of the zymogen (pro ADC). The class I of ADC cleavage requires the acetyl-CoA sensor MRF (Maduration Regulatory Factor) and has been found only in gamma proteobacteria. Class II of ADCs is an autocatalytic cleavage and is found in a wide number of bacterial phyla. Since the majority of archaea lack homologs of the *E. coli* acetyl-CoA synthesis pathway genes, the mechanism of pantothenate/CoA biosynthesis has not been completely deduced in this organism. This pathway includes a glutamate decarboxylase (GAD) that substitutes the ADC and takes pyridoxal 5’-phosphate (PLP) as a cofactor, was reported in archaea (5). Curiously, GAD prefers aspartate (Asp) as opposed to glutamate (Glu) as its substrate although, commonly GAD catalyzes the decarboxylation of Glu to γ-aminobutyrate (GABA).

Although prokaryotes and eukaryotes have an indispensable requirement for β-ala for the synthesis of coenzyme A (CoA), the pathways involved in its synthesis are very diverse. The uracil fermenting bacterium *Clostridium uracilicum* degrades uracil to β-ala. Uracil or thymine is first converted to dihydrouracil. The enzyme dihydropyrimidinase catalyzes the hydration of dihydrouracil to produce *N*-carbamoyl- β-ala, which is hydrolyzed to β-alanine, CO_2_ and NH_3_ by β-ala synthase (6).

The reductive degradation of pyrimidine as a source of β-ala was supported by genetics and biochemical analysis in several bacteria including *Clostridium uracilicum* (6) and *Clostridium botulinum* (7). Although the reductive degradation of pyrimidines has also been implicated as the *de novo* source of β-ala in *E. coli* auxotrophs, lack of response to dihydrouracil indicated that in these bacteria the major pathway for β-ala synthesis was the decarboxylation of aspartate catalyzed by ADC.

Genschel (8) performed a phyletic distribution of *E. coli* and human genes involved in pantothenate and CoA synthesis across 47 completely sequenced genomes of 20 Bacteria, 16 Archaea, and 11 Eukaryotes. This study revealed a mosaic of orthologs with 20 to 70 % amino acid identities. At least, one protein was missing from each of the 47 genomes analyzed.

Comparative genomics using the *E. coli* pantothenate pathway genes as query across twenty sequenced bacterial genomes revealed multiple gaps that may represent distantly related homologues due to the absence of at least, one gene per bacterial genome surveyed (8).

The order *Rhizobiales* is a heterogeneous group of Gram-negative bacteria, located taxonomically within the division of alpha proteobacteria. Some of its members are facultative diazotrophs that associate with leguminous plants to carry out symbiotic nitrogen fixation. Others are pathogens of plants or animals (9). Our model is *Rhizobium etli* CFN42, isolated from bean root nodules (10). Its genome consists of a circular chromosome and six large plasmids ranging in size from 194 to 642 Kb (11).

In the course of examining *Rhizobium etli* CFN42 plasmids for the presence of housekeeping genes encoding essential functions, we found that both *panC* and *panB* genes were clustered together on the 642 kb replicon p42f. We demonstrated that both are indispensable for the synthesis of pantothenate (12) (Fig. S1). Surprisingly, we did not find homologues of the *E. coli panD* gene in the genome of *R. etli* CFN42. Since strain CFN42 grows in minimal medium without pantothenate or β-ala addition, it was assumed that it is a pantothenate prototroph. *Agrobacterium fabrum* C58 (formerly *A. tumefaciens* C58), a plant pathogen that induces tumors in numerous plants, was the only member of *Rhizobiales* order included in Genschel’s study. According to this analysis, *A. fabrum C58* lacks ketopantoate reductase (KPR, EC 1.1.1.1.169) but has a putative ADC detected by a BlastP search. We performed BlastP searches in order to get insight on the presence or absence of ADC in the genomes of rhizobial reference strains.

Several questions arise from the presence or absence of ADC in *R. etli* CFN42 and *A. fabrum*. Is the absence of ADC an exclusive characteristic of strain CFN42, or is it a widespread characteristic of the *Rhizobiales* order or perhaps the α-proteobacteria?

The aim of this work was to identify the biosynthetic origin of β-ala that replaces the function of ADC allowing *R. etli* CFN42 to be a β-ala prototroph. We also performed an *in-silico* analysis of the α-proteobacteria group to understand the occurrence, diversity and evolution of the enzymes involved in β-ala synthesis.

## Results

### Orthologues of the canonical L-aspartate-α-decarboxylase enzyme are predominantly absent in α-proteobacteria

A previous study on ADC phylogeny and amino acid conservation analyses revealed that ADCs are in γ-proteobacterial genomes and most of them maintain the *panCBD* synteny (4). We noticed the absence of the *panCBD* gene cluster while studying the functional characterization of *panC* and *panB* in rhizobia (12). In the present study, BlastP and Psi Blast searches using as query ADC from *E. coli* and *A. fabrum* C58 revealed the absence of ADC homologues in *R. etli* CFN42 and other reference strains.

To generalize the absence of ADC homologues in α-proteobacteria, we assessed the occurrence of putative ADCs in the proteome of 204 α-proteobacteria (84 rhizobia and 120 members of seven families of α-proteobacteria (Table S1). The complete proteome of each bacterium was obtained from the NCBI reference sequence collection (RefSeq) and clustered with ProteinOrtho v5.15 a large-scale Blast-based orthology detection tool (13) (Fig. S2). This analysis only showed 37 putative ADC from 204 α-proteobacteria genomes.

### Unrooted ML-based tree inferred from the α- and γ-proteobacteria ADCs revealed high divergence among them

An important characteristic of the α-proteobacteria is its genome plasticity that allows different genome rearrangements including deletions or duplications (14, 15). We made a phylogenetic analysis to get a wider view on the evolutionary relationship among the ADCs from α- and γ-proteobacteria This ML-based tree inferred from the α- and γ-proteobacteria ADCs is shown in fig. 1, (Table S2). To determine if this ADC phylogeny maintains the coherence of species phylogeny, it was compared to the previously reported species trees performed by the Bayesian analysis of 104 concatenated alignments (16); and with the most recent robust species tree, this was done under the maximum likelihood framework with a data set of 200 single-copy and conserved genes for the α-proteobacteria (17).

**Figure 1.**
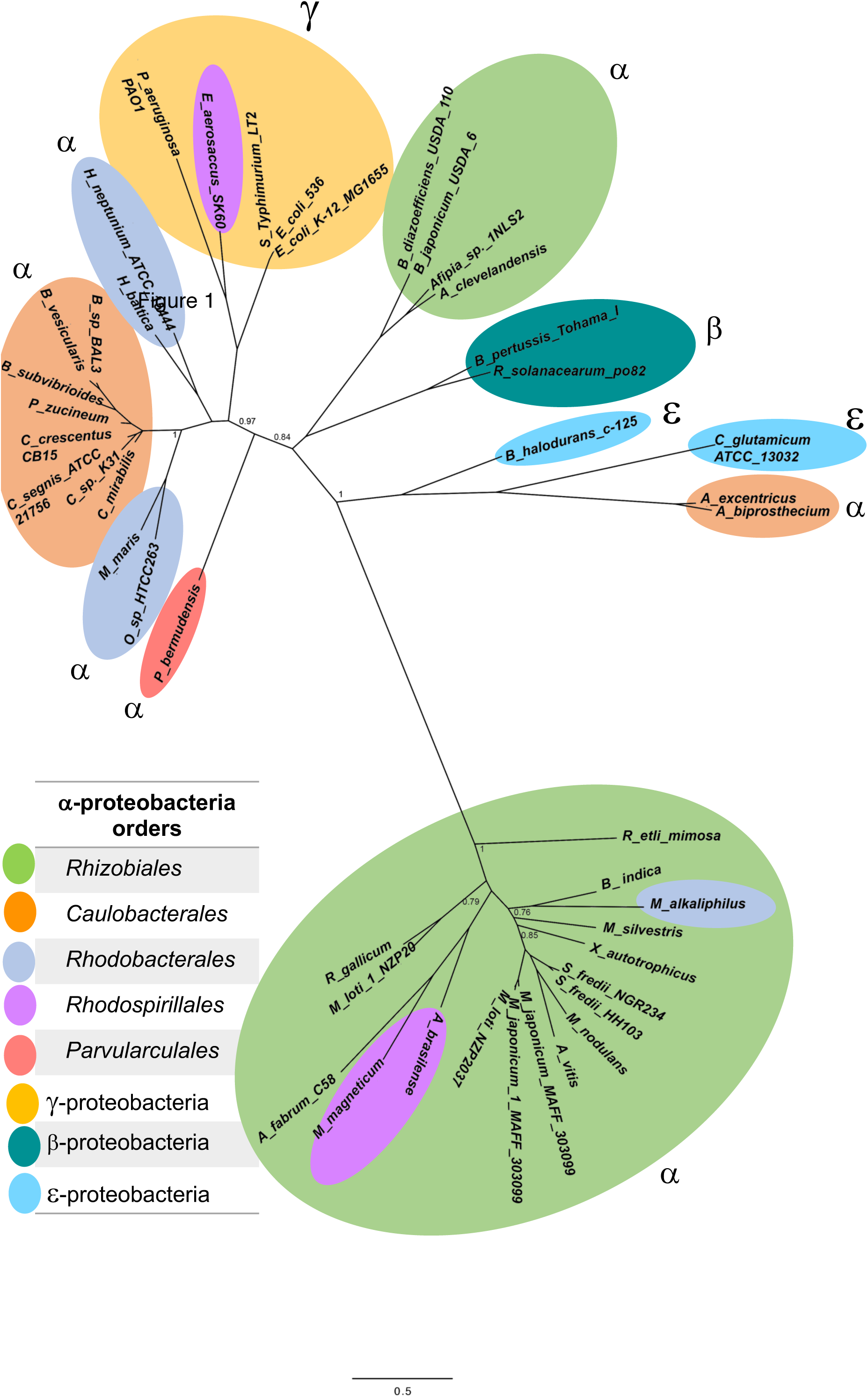
The putative ADCs of α-proteobacteria found in our occurrence analysis. A maximum likelihood phylogenetic tree inferred from a subset of 204 genomes, where we extracted only 37 ADCs. The tree shows a monophyletic clade of proteins distantly related to those from γ-proteobacteria and other α-proteobacteria.

The majority of ADCs belonging to the order *Rhizobiales* were grouped in a sole cluster (Fig. 1, green oval bottom). Unexpectedly, we found two ADCs from *Bradyrhizobium japonicum* and *Afipia sp* close to γ- and β-proteobacteria (Fig. 1 upper green oval). Two ADC of the *Rhodospirillales Order, (Azospirilum brasilense and Magnetospirillum magneticum)*, were located as orthologs of *Rhizobiales* (Fig. 1, purple oval). The ADC from *Maritimibacter alkaliphilus* was located within the order *Rhizobiales* whereas in the species tree *M. alkaliphilus* belongs to the *Rhodobacterales* order (17).

This heterogeneous cluster of *Rhizobiales* ADCs links through a long branch with remote orthologues from class α-proteobacteria belonging to Orders: *Rhizobiales* (*B. japonicum* and *Afipia) Caulobacterales* (C. crecentus), *Rhodobacterales* (*Hyphomonas neptuniou* and *Hirschia baltica*), *Parvularculales* (Parvacula bermudensis γ-proteobacteria (outgroup, *E. coli, Salmonella, Pseudomonas aeruginosa*), β-proteobacteria (*B. pertussis* and *R. solanacearum*), ε-proteobacteria (*Corynebacterium glutamicum*).

We observed the clustering of 14 sequences found on the *Rhizobiales* clade, including two exceptions from *Rhodobacteriales* order and one from *Magnetoccocales* order. The *Rhizobiales* order and the rest of α-proteobacteria, are clearly apart, this correlates with the loss on amino acid conservation, suggesting an event of divergence on ADC *Rhizobiales* homologs toward proteobacteria ones.

### Presence, absence, duplications and functional redundancy of the six genes involved in synthesis of pantothenate

In addition to ADC, the 84 rhizobial genomes were surveyed for the presence of the enzymes that catalyze the pantothenate synthesis. This study revealed that the enzyme KPHMT (ketopantoate hydroxymethyl transferase, PanB) is highly conserved in the *Rhizobiales* order and was absent in only 8.4% of the genomes analyzed. The genera lacking KPHMT were: *Bradyrhizobium* sp ORS_278, *Candidatus_liberobacter* (4 strains), and *Hoeflea* (2 species). KPHMT was predicted to be present in the other members of *Rhizobiales*, which have a diversity of habitats. Two copies of this enzyme were present in 17.8% of rhizobia, mostly in *Rhizobium* and *Sinorhizobium* species. Three copies of KPHMT were found in 4.8% of the *Rhizobiales* order: three *Mesorhizobium* species and one in *Rhizobium leucaenae.*

The next step on the pathway is the reduction of α-ketopantoate to produce pantoate. Two enzymes can perform this reduction, KPR (α-ketopantoate reductase (PanE) was found in only 57% of the Rhizobial genomes while KAR (acetohydroxy acid reductoisomerase, ilvC) was present in 95.2% of the genomes. Most of human, plant and mammalian pathogens have lost the KPR enzyme. Interestingly *Candidatus* genera lacked both KAR and KPR enzymes in their genomes.

In the last step of the pathway, pantothenate synthetase (PS, PanC) catalyzes the ATP-dependent condensation of D-pantoate with β-ala to form pantothenate. This enzyme was absent in 7.1% of the genomes surveyed, some of which belong to parasites such as *Hoeflea* and *Candidatus.* Strains with a single copy were found in 88% of the rhizobia genomes analyzed. Gene duplications were found in 4.7% on *Mesorhizobium* and *Bradyrhizobium* species.

The occurrence of putative ADC enzymes is shown in Table 1. An ADC encoding gene was present in 19 % of the genomes, and 2 strains had a second copy of ADC in their genomes. *Mezorhizobium japonicum* MAFF303099 had one in chromosome and the other one in a plasmid, and M. loti NZP 2037 had both in chromosome.

**Table 1.** Ocurrence of pantothenate synthesis genes on *Rhizobiales* order

| *Gene* | *Enzyme* | *Occurrence* |
| --- | --- | --- |
| *panD* | ADC | 19.04% |
| *panM* | MRF | 21.42% |
| *panB* | KPHMT | 91.66% |
| *panE* | KPR | 57.14% |
| *ilvC* | KAR | 95.23% |
| *panC* | PS | 92.85% |

The search for MRF (Maturation Regulatory Factor, *panM*) homologues revealed that 78.5% of the genomes lacked an MRF homologue; 15.4% had one copy and 6% encoded two copies. However, only five genomes coded for both ADC and MRF *(A. fabrum* C58, *Bradirhizobium japonicum* USDA11, *R. etli bv mimosa* str. Mim1, *Rhizobium gallicum*, and *Sinorhizobium fredii* HH130).

Our results showed that only 16 of the 84 genomes analyzed (19.04%), encoded the complete pantothenate pathway. Of the 68 genomes with gaps in the pantothenate pathway, the predominant deficiencies were a lack of ADC in 80.95 % of the genomes and the absence of both ADC and KPR in 38%.

### *R. etli* CFN42 is a pantothenate prototroph

The model of pantothenate synthesis established in *E. coli* (1, 3), indicates that the enzymes missing in rhizobia should cause auxotrophy. Growth assays were done in liquid minimal medium of the wild type strains *R. etli* CFN42 (lacks *panD*) and *Sinorhizobium melilloti* 1021 (lacks *panD* and *panE*), and a *R. elti* CFN42 plasmid p42f cured strain (CFNX186) that is defective for growth in minimal medium without pantothenate. We found that the wild type strains were able to grow after three sub-cultures in minimum medium without β-ala or pantothenate. This shows that even with the absence of panE and/or panD rhizobia are still able to synthesize β-ala and pantothenate (Fig.2). This prototrophy contrasts with the auxotrophy exhibited by *R. etli* CFNX186, which lacks *panC* and *panB* as well as plasmid p42f (18).

**Figure 2.**
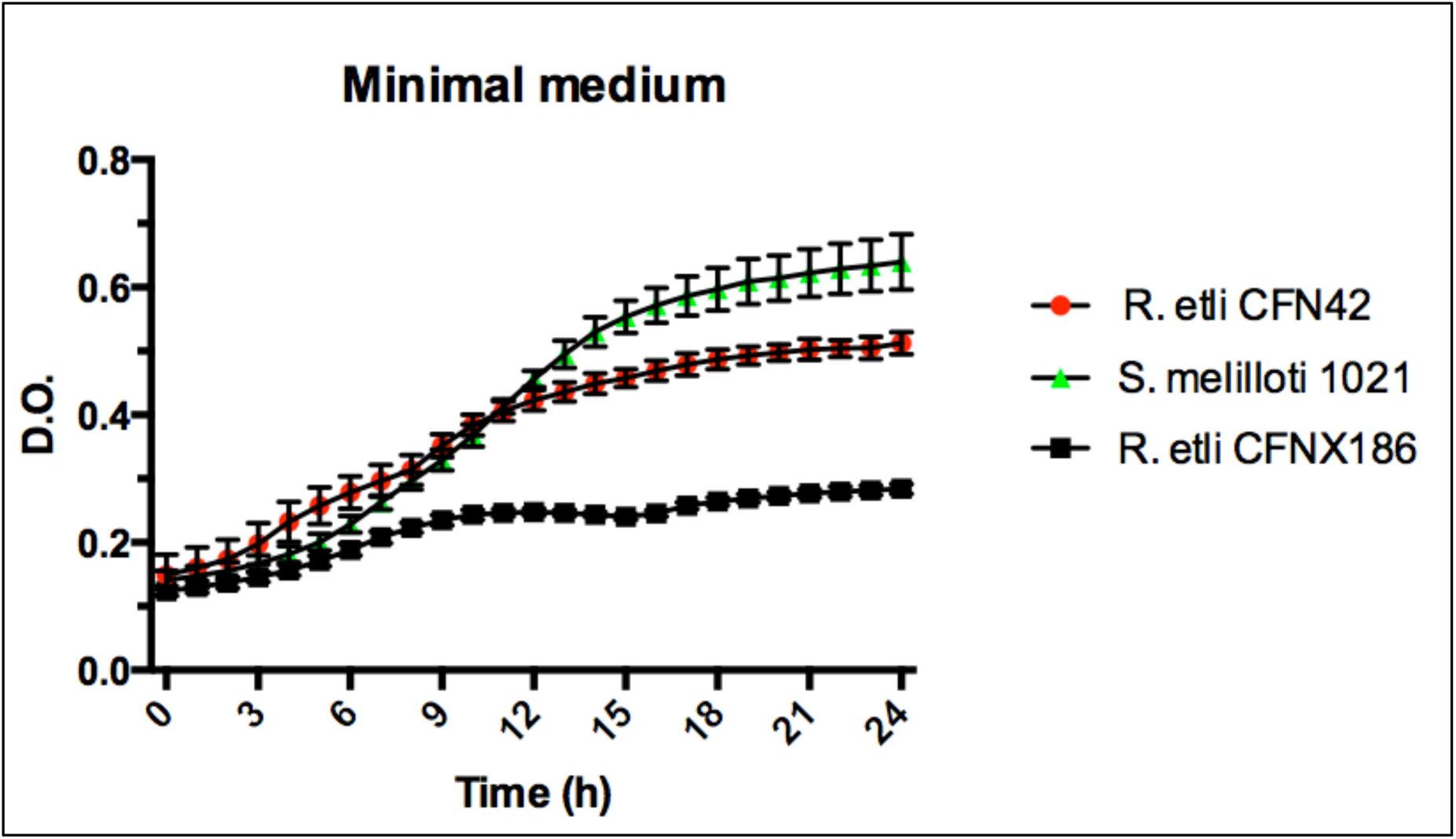
Growth test for prototrophy of wild type *R. etli* CFN42, its p42f-cured derivative CFNX186, and in wild type *S. meliloti* 1021. Tested in minimal medium without β-alanine or pantothenate.

### Occurrence of different pathways that would replace ADC in rhizobia

To identify which enzyme(s) might be responsible for the synthesis of β-ala, we performed bioinformatic analyses of 204 alpha proteobacterial genomes to find possible pathways or genes that could potentially produce this metabolite. Based on a literature search we selected 6 genes of interest that encode enzymes of the pyrimidine degradation pathway (AmaB, Dht, PyrD), glutamate decarboxylase (GAD) and the transaminases AAM and GabT (Fig. 3).

**Fig. 3.**
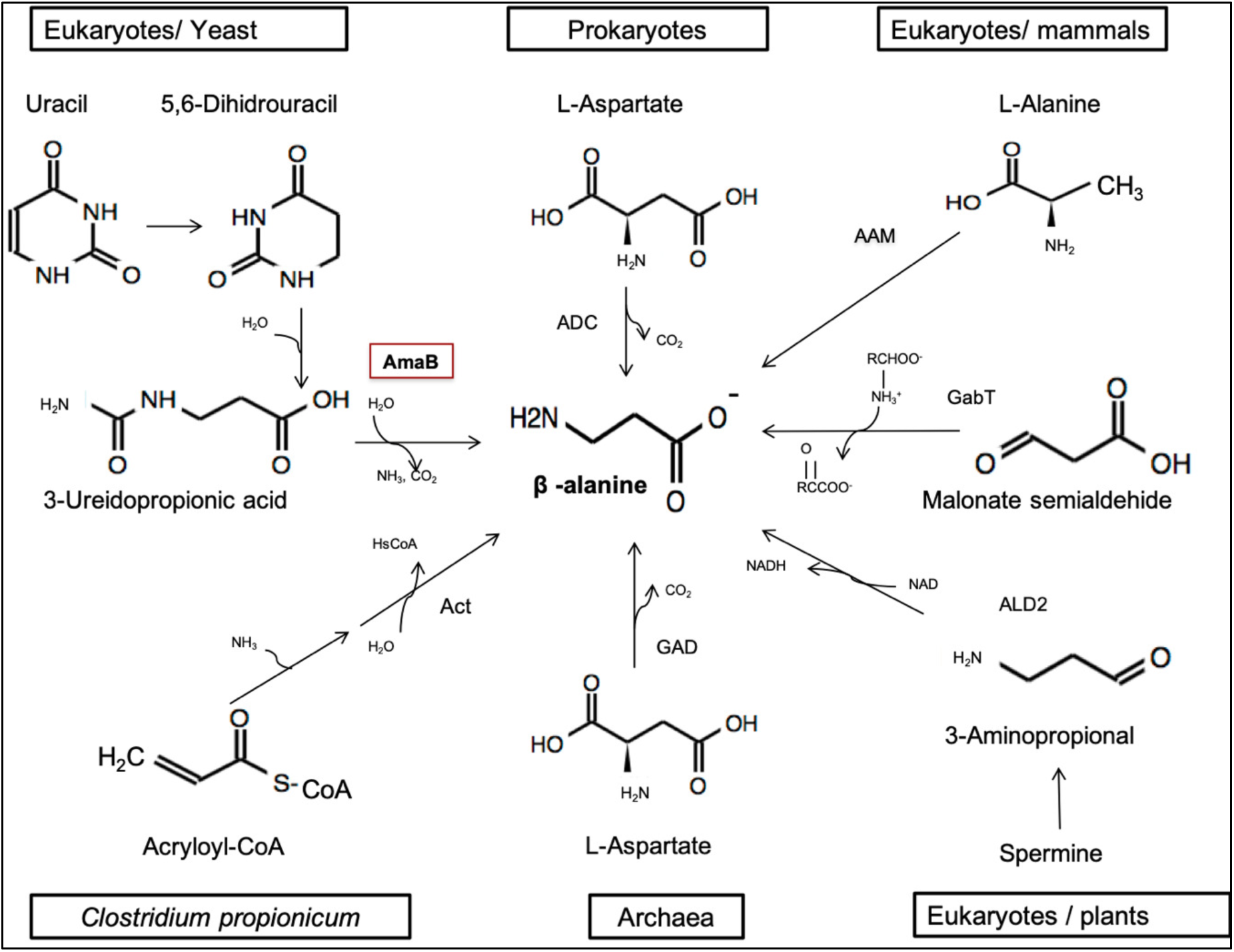
β-alanine biosynthesis in different domains of life. (AmaB) β-alanine synthase; (ADC) 1-Aspartate decarboxylase; (AAM) 2,3-aminomutase; (gabT) 4-aminobutirate transferase; (ALD2) amine oxidase; (GAD) glutamate decarboxylase; (Act) β-alanyl-CoA:ammonia lyase.

It is believed that, β-ala synthesis in bacteria results only through decarboxylation of aspartate by ADC (19, 20). Other ways of producing β-ala exist in the other domains of life. Two routes occur in fungi: *Saccharomyces cerevisiae* produces β-ala by the degradation of spermine (21), and *Schizosaccharomyces pombe* and *Saccharomyces kluyveri* obtain it from uracil degradation (22).

The pyrimidine degradation pathway involves three enzymatic steps from uracil to produce β-ala, CO_2_ and NH_3_ (6). In the final step of the pathway, β-ala synthase (AmaB) uses N-carbamoil-β-alanine as substrate. In rhizobia, the *in vitro* activity of AmaB has been detected in *A. fabrum* C58 and *S. melilloti* 1021. The authors showed the production of β-ala from 3-ureidopropionic acid *in vitro*, in the last step of the pathway (23).

In archaea, β-ala can be synthesized by a GAD that uses Asp as a substrate. In these studies, it was shown that two enzymes annotated as GADs had higher affinity for Asp than for Glu and, they demonstrated the *in vitro* activity of the enzymes in *Methanocaldococcus jannashi* and *Termococcus kodakarensis* (5, 24).

We also included in the study two transaminases that in bacteria, insects and mammals produce β-ala in a single-step reaction. The first one, AAM acts on L-alanine and 3-oxopropanoate to produce pyruvate and β-ala (25, 26). The second, GabT performs a transamination of malonate semialdehyde and L-glutamate (27, 28).

In summary, our bioinformatics analysis showed that two transaminases and the pyrimidine degradation pathway were encoded in the *R. etli* CFN42 genome. We did not find any candidate genes for ADC or GAD, or any complete polyamine degradation pathway.

### AmaB functionally complement ADC loss

In our study we tested the function of different genes in *R. etli*, by inactivating those that encode two transaminases (AAM and GabT) and the *amaB* gene for pyrimidine degradation (Table S3). Following with the canonical decarboxylation pathway, we found a putative ω-amino acids decarboxylase that was different from the ADC and GAD enzymes. The genes were interrupted using a suicide plasmid, and the resulting mutants tested for growth in minimal medium without β-ala or pantothenate. From this screening we found that the *amaB* mutant was auxotrophic for β-ala, while inactivation of the other genes caused no growth deficiency (data not shown).

AmaB (RHE_CH03290) is a chromosomal gene annotated as β-alanine synthase. It belongs to the pyrimidine degradation pathway and transforms 3-ureidopropionic acid to β-ala, CO_2_ and ammonia. We disrupted this gene in *R. etli* CFN42 and grew the mutant ReAM-1 (*amaB*^*-*^) in minimal medium (MM) without β-ala or pantothenic acid. The mutant was deficient in growth, indicating a β-ala auxotrophy, and its growth was restored by exogenous β-ala or by introducing the *amaB* gene on a plasmid (Fig. 4).

**Fig. 4.**
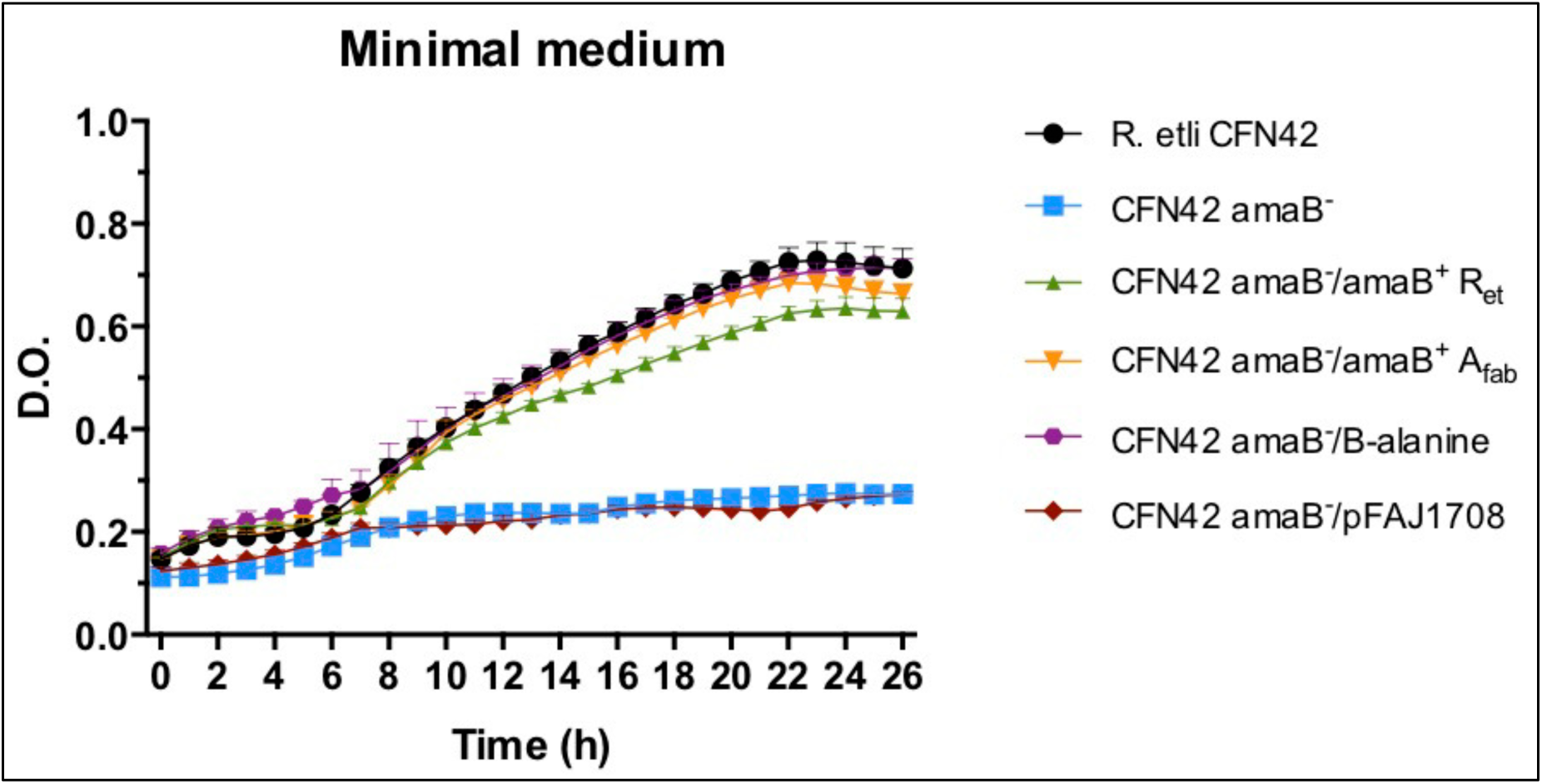
Growth in minimal medium of the *R. etli* CFN42 wild-type (•) strain and its derivative complements. *R. etli* CFN42 *amaB* mutant (▪); CFN42 *amaB*^*-*^ */amaB*^*+*^ of *R. etli* CFN42 (•); CFN42 *amaB*^*-*^*/amaB*^*+*^ of *A. fabrum* C58 (•); CFN42 *amaB*^*-*^ complemented with 1uM of β-alanine (•); CFN42 *amaB*/pFAJ1708 empty vector (•).

Similarly, the mutant was complemented with a plasmid-borne copy of the *amaB* gene from *A. fabrum* C58. The product of this gene has been shown to have β-alanine synthase activity *in vitro* (29).

### Purified AmaB produces β-ala from 3-ureidopropionic acid *in vitro*

The his_6_-tag enzyme was purified on an immobilized nickel affinity column under native conditions and had a molecular mass of 60 kDa, consistent with β-ala synthase (45 kDa) plus the 15 kDa 6His-Sumo tag (Fig. S3).

Enzymes of this type are characterized as metallo-enzymes that use Ni^2+^ and Co^2+^ as cofactors in enzyme assays. The reaction mixture contained purified AmaB pre-incubated with Ni^2+^ or Co^2+^, 10mM MgCl_2_, 100mM sodium phosphate buffer and 3-ureidopropionic acid as a substrate. We initially used a TLC system with ninhydrin detection to identify the present of β-ala (30) (Fig. S4). We observed enzymatic activity with both metal ions and no product was formed in their absence.

As described below, we also performed our enzymatic assays using an HPLC system to obtain a better resolution.

### Synthesis of β-alanine by recombinant AmaB

AmaB was heterologously expressed in *E. coli* strain BL21 and recovered by Ni^2+^ affinity chromatography as previously described (29). Production of β-ala by recombinant AmaB was analyzed by HPLC. The fluorescence response of a β-ala was linear with concentration (Fig. S5). The time-course of recombinant AmaB activity using 3-ureidopropionic acid as substrate and Ni^2+^ as cofactor, showed that β-ala is synthesized at a linear rate for up to 30 min (Fig. 5A). β-ala was not detected in a reaction assay without recombinant AmaB protein (Fig. 5B). The standard of β-ala overlapped with the peak of the compound synthesized by AmaB, while the α-ala standard did not (Fig. 5C). These results indicated that recombinant AmaB is able to synthesize β-ala.

**Fig. 5.**
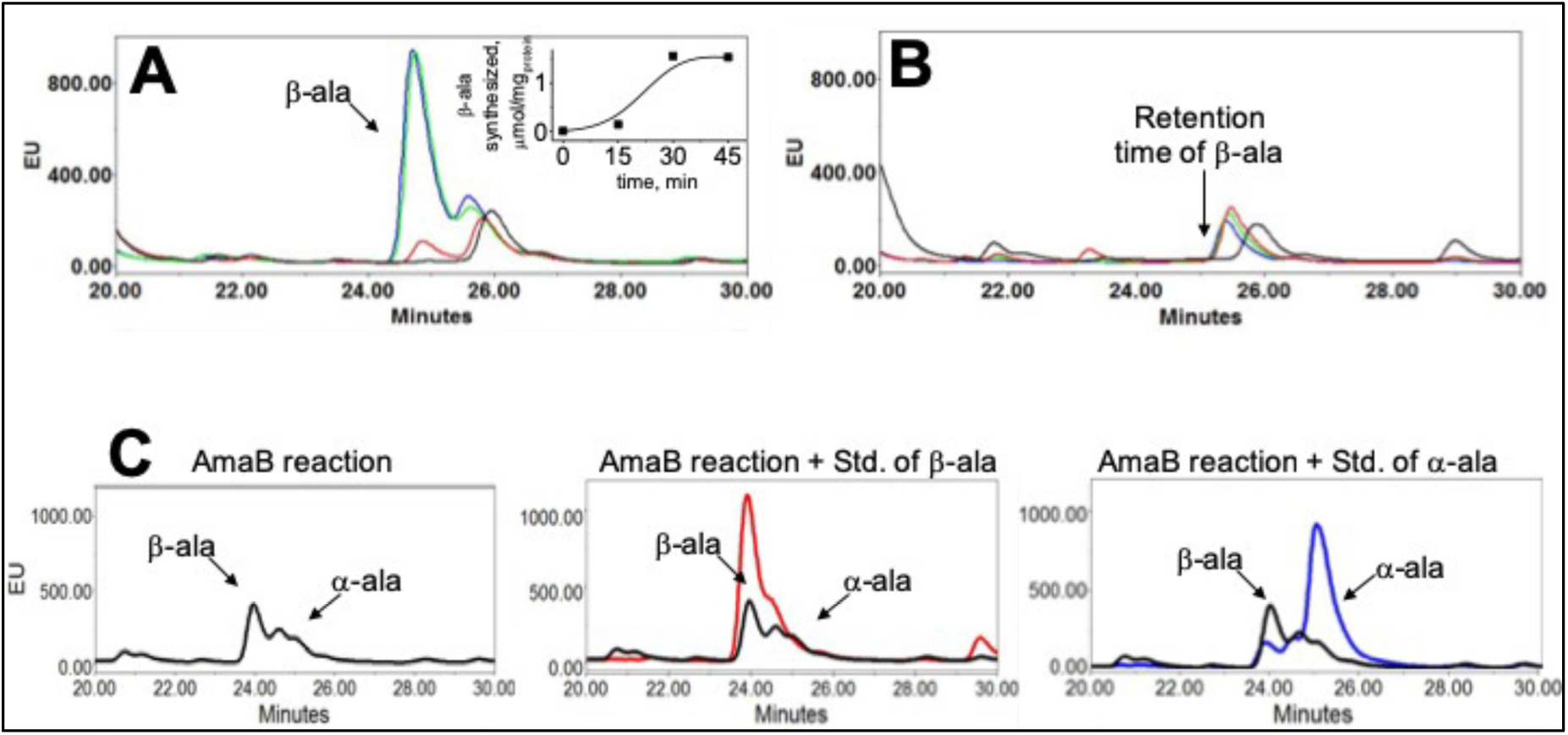
Analysis β–alanine synthesized by AmaB. (A) Activity of recombinant AmaB, at 0 (—), 15 (—), 30 (—) and 45 min (—). Peak of β-ala was observed at 24.8 min. (B) Reaction without enzyme, β-ala peak was not observed. (C) To confirm peak of β-ala, internal standard of β-ala (—) and L-ala (—) were added independently to neutralized AmaB reaction, carried out for 15 min (—).

## Discussion

The relevance of β-ala as a key component of pantothenate synthesis has been well established. However, the synthesis of β-ala had not been totally elucidated. Pioneer studies performed in *E. coli* and γ-proteobacteria defined that β-ala was synthesized by the decarboxylation of L-aspartate in a one-step reaction catalyzed by ADC. The concept of a canonical one-step decarboxylation reaction prevalence was assumed to be the sole source of β-ala in bacteria for many years.

The genomic era facilitates pathways comparison among numerous species (8), this bioinformatic approach helped us to determine the diversity of mechanisms involved in β-ala synthesis. In this study, we found several differences between *R. etli* (α-proteobacteria) and *E. coli* (γ-proteobacteria) the most intriguing was the absence of an ADC homolog in Rhizobia. In previous studies, analyses of *E. coli* and other γ-proteobacteria revealed that β-ala was produced by the decarboxylation of aspartate by aspartate decarboxylase enzyme (ADC) (20); whereas in several Archaea β-ala was the synthetized by a glutamate decarboxylase (GAD) able to decarboxylate both aspartate and glutamate (5). These data confirm the relevance of one-step decarboxylases not only in bacteria but also in Archaea.

An alternative source of β-ala synthesis is the reductive degradation of pyrimidine. This three-step reaction was found in: *Clostridium uracilicum* (6), *C. botulinum* (7), as well as in *E. coli* strains: *E. coli* W, *E. coli* D2, *E. coli* 99-1, and E. coli 99-2 (Table S1), that in contrast with previous studies none of them was able to growth in the presence of dihydrouracil and β-ureidopropionic acid (31).

In bacteria belonging to the order *Rhizobiales* little is known about the metabolism of β-ala and pantothenate (12). The occurrence analysis performed in in this work indicates that our model, *R. etli* CFN42 lacks ADC and GAD the most common one-step reaction used in bacteria to synthesize β-ala. We suggest that there can be functional redundancy on certain rhizobia strains. As part of our work, we constructed different single and double mutants on *A. fabrum* C58, to try to get an auxotrophic strain, but in all cases the mutants were prototrophic (data not show).

Particularly for the *Rhizobiales* order, we constructed a heat map with their most representative genomes; here we can associate the loss and prevalence of different pathways, assuming that the decarboxylation pathway is missing on most of rhizobia genomes (figure S6).

*S. meliloti* and *A. fabrum C58*, have tested for production of β-amino acids through the uracil degradation pathway because of their pharmaceutical relevance (23). Unexpectedly, the research only showed the ability to produce β and ω amino acids *in vitro*; we do not know if these strains synthesize β-ala through this pathway or if these strains have a functional redundancy with another β-ala synthesis pathway.

As part of our occurrence analysis, we extended our work to alpha-proteobacteria with 120 more genomes from seven different orders, (Table S1). We found a correlation between the rhizobia order and alpha proteobacteria. In general, we observed that the pyrimidine degradation pathway (37%) and AAM transaminase (56%) are widely distributed in alpha proteobacteria, as well as in the rhizobia order (Table S1). Also, we observed that ADC and GAD enzyme are poorly represented in alpha-proteobacteria, with 17% and 6.8% respectively. This analysis suggests a strong co-relation between the decarboxylation pathway loss and predominance of pyrimidine degradation pathway in the *Rhizobiales* order and α-proteobacteria.

Also, we tested the activity of recombinant AmaB *in vitro* by HPLC, to confirm the catalytic activity of this protein to produce β-ala from 3-ureidopropionic acid corroborating the presence of a different way in bacteria to produce this essential metabolite.

## MATERIALS AND METHODS MATERIALS AND METHODS

### DNA manipulations

Standard techniques were used for plasmid and total DNA isolation, restriction, cloning, transformations, and agarose gel electrophoresis (32). Plasmid mobilization from *E. coli* to *Rhizobium* was done by conjugation performed on PY plates at 30°C by using overnight cultures grown to stationary phase. Donors (*E. coli* DH5) and recipients (*R. etli* CFN42 wild type and mutant strains) were mixed at a 1:2 ratio, and suitable markers were used for transconjugant selection.

### Bacterial strains, media and growth conditions

The characteristics of the bacterial strains and plasmids used in this study are listed in Table S3. Bacterial growth was started from glycerol stocks (20% and stored at -70°C) propagated in rich PY medium (per L, 5 g peptone, 3g yeast extract, and 15g agar). Rhizobium strains were grown at 30°C in three different media: a) PY rich medium, b) Minimal medium (MM) and c) Minimal medium plus 1 μM calcium pantothenate (MMP). MM was prepared as follows: a solution containing 10 mM succinate as carbon source, 10 mM NH_4_Cl as nitrogen source, 1.26 mM K_2_HPO_4_, 0.83 mM MgSO_4_, was adjusted to pH 6.8 and sterilized. After sterilization the following components were added to the final concentration indicated: 0.0184 mM FeCl_3._6H_2_O (filter sterilized), 1.49 mM CaCl_2.2_H_2_O (autoclaved separately), 10μg/ml biotin and 10μg/ml thiamine (both sterilized by filtration). MMP contains the same components plus 1 μM calcium pantothenate.

The rhizobia strains were cultivated at 30°C for 20h in PY medium. *E. coli* K12 MG1655 and *Escherichia coli* BL21 was used to clone and express the β-alanine synthase gene. It was cultivated at 37°C for 20h in Luria-Bertani medium (LB medium; 1% tryptone, 0.5% yeast extract, 0.5% NaCl, pH 7.2).

### Cloning and sequence analysis of *amaB* gene, mutants and complementation

The *ama*B (RHE_CH03290) gene was overexpressed in *E. Coli* DH5aalpha. The coding region of the *amaB* gene was amplified from the genomic DNA of *R. etli* CFN42 by PCR using primer set AmaBF1 (5’ATGGTGGCAGCACCAGGCGAGAACATGC-3’); AmabR1 (5’-TCACACCACGATCTCCGCCGTCTCCACC-3’). The amplified fragment was inserted into the pET-SUMO expression vector (Ni-NTA Purification System sigma Aldrich). After confirming the absence of unintended mutations, the plasmid was introduced into *E. coli* strain BL21(DE3). Primer sets list are on table S4.

### Overexpression and purification of wild-type β-ureidopropionase AmaBret

The transformant BL21 strain was grown in LB medium supplemented with 100 mg/ml^□^ of carbenicilin. A single colony was transferred into 10 ml of LB medium with carbenicilline at the above-mentioned concentration in a 100-ml flask. This culture was incubated overnight at 37°C with shaking. Five hundred milliliters of LB medium with the appropriate concentration of carbenicilin was inoculated with 5 ml of the overnight culture in a 1-liter flask. After 3 h of incubation at 37°C with vigorous shaking, the optical density at 600 nm (OD_600_) of the resulting culture was 0.3 to 0.5. For induction of β-alanine synthase gene expression, isopropyl-β-D- thiogalactopyranoside (IPTG) was added to a final concentration of 0.1 mM, and incubation was continued at 30°C for an additional 6 h.

The cells were collected by centrifugation (8,000 x *g*, 4°C, 10 min), washed twice, and re-suspended in 50 ml wash buffer (2.5 M NaCl, 250mM NaH_2_PO_4_, 20mM imidazole pH 8.0). The cell walls were disrupted by sonication using a UP 200 S ultrasonic processor, on ice for fourth periods of 15 s at pulse mode 0.5 and 40% sonic power. The cell debris was precipitated by centrifugation (Sigma 12169-H rotor; 8,000 x *g*, 4°C, 10 min), and the supernatant was applied to a column with 2ml ml of Niquel metal-affinity resin (Ni-NTA Purification System sigma Aldrich). After being washed four times with wash buffer, B-Ureidopropionase enzyme was eluted with elution buffer (500 mM NaCl, 3M imidazole, 20 mM Sodium phosphate buffer, pH 6.0). The purified enzyme was dialyzed against 20 mM sodium phosphate buffer, pH 8.0, and stored at 4°C until use.

### Enzyme assays

The standard enzymatic reaction was carried out with purified **AmaB_ret_** (at a final concentration of 1mg/ml) along with 125 mM 3-Ureidopropionic acid and 10 mM MgCl_2_ dissolved in 100mM sodium phosphate buffer pH 8.0, in a 3ml reaction volume (29). The reaction mixture was incubated at 30°C for 60 min, with the apoenzyme preincubated (1 h) at 4°C with 2mM of NiCl_2_ and 500ul aliquots were stopped each 15 minutes, by the addition of 50ul of 3% TCA. After centrifugation, the resulting supernatants were analyzed by high-performance liquid chromatography (HPLC).

### Determination of β-Alanine by HPLC-florescence

Determination of β-alanine (β-Ala) synthesis was carried out by HPLC coupled to a Multi λ–fluorescence detector (Waters 1525/2475, Milford, MA, USA) using a reverse-phase C-18 Spherisorb ODS2 column of 5 μm particle size and 150 × 4.6 mm (Waters, Mildford, USA) (33). Enzymatic reactions were stopped with perchloric acid (3% v/v) at indicated times and immediately frozen in liquid nitrogen and kept at - 70C°. The acidic supernatants were neutralized with 3 M KOH/0.1 M Tris and centrifugated to remove KClO_4_. Supernatant were recovered and used for β-Ala determination by derivatization with 37 mM ortho-phthalaldehyde (OPA). The β-Ala and α-Alanine standards (Sigma-Aldrich, Saint Louis, MO, USA) were used for identifying of chromatographic peaks.

### Identification of orthologous proteins with multiple sequence alignments and phylogenetic analysis

We selected 204 alpha-proteobacteria organisms to analyze the presence and absence of 12 proteins related to the pantothenate synthesis and transport. The protein FASTA files (faa) for each of the 204 alpha-proteobacteria genomes were downloaded from the RefSeq NCBI database. Protein sequences with an expectation value (E) of 10^-3^ or less were considered as putative homologues. We used Proteinortho v5.15 to obtain the clusters of orthologous proteins from the 204 protein FASTA files. Next, we used the Pfam v31.0 database to determine which proteinortho clusters represent the 12 proteins of interest used in this work. The proteins we searched for were: PYD1, PYD2, PYD3, GAD, KPHMT, PS, ADC, KPR, MRF, KAR, AAM and GabT. Finally, we determined which alpha-proteobacteria organisms were represented in each protein cluster.

For phylogenetic analysis, we used the Pfam v31.0 database to determine which proteinortho clusters represent the ADC proteins. A total of 37 homologs belonging to alpha proteobaceria sequences were tested with a group of nine external sequences listed in Table S2 and, were aligned against Muscle v3.8.31 (30). The resultant data set containing 46 putative ADC’s homologs, was used to infer the evolutionary relationships. We used ProtTest3 v3.4.2 for the evolutionary model and, the best result was LG+G model; using amino acid alignment. The phylogenetic analysis was performed with PhyML v3.3.20170530 under (-d aa -m LG -a e -o ltr) parameters (31).

## ACKNOWLEDGMENTS

Mariana López-Sámano is deeply grateful to Programa de Doctorado en Ciencias Biomédicas belonging to Universidad Nacional Autónoma de México (UNAM), where she conclude her doctoral studies. She also acknowledge to CONACYT for the fellowship received. We are grateful to Susana Brom and Michael F. Dunn for helpful discussion and critically reviewing the manuscript. We gratefully acknowledge to Laura Cervantes and Victor Antonio Becerra Rivera for their skillful technical assistance.

